# Moving microbes: the dynamics of transient microbial residence on human skin

**DOI:** 10.1101/586008

**Authors:** Roo Vandegrift, Ashkaan K. Fahimipour, Mario Muscarella, Ashley C. Bateman, Kevin Van Den Wymelenberg, Brendan J.M. Bohannan

## Abstract

The human skin microbiome interacts intimately with human health, yet the drivers of skin microbiome composition and diversity are not well-understood. The composition of the skin microbiome has been characterized as both highly variable and relatively stable, depending on the time scale under consideration, and it is not clear what role contact with environmental sources of microbes plays in this variability. We experimentally mimicked human skin contact with two common environmental sources of microorganisms — soils and plant leaves — and characterized the dynamics of microbial acquisition and persistence on skin on very short time scales. Repeatable changes in skin community composition following encounters with environmental sources were observed, and these trajectories largely depend on donor community biomass distributions. Changes in composition persisted for at least 24 hours and through a soap and water wash following exposures to relatively high biomass soil communities. In contrast, exposures to lower biomass leaf communities were undetectable after a 24 hour period. Absolute abundances of bacterial taxa in source communities predicted transmission probabilities and residence times, independent of phylogenetic considerations. Our results suggest that variability in the composition of the skin microbiome can be driven by transient encounters with common environmental sources, and that these relatively transient effects can persist when the source is of sufficient biomass.

**Importance:** Humans come into contact with environmental sources of microbes, such as soil or plants, constantly. Those microbial exposures have been linked to health through training and modulation of the immune system. While much is known about the human skin microbiome, the short term dynamics after a contact event, such as touching soil, have not been well characterized. In this study, we examine what happens after such a contact event, describing trends in microbial transmission to and persistence on the skin. Additionally, we use computational sampling model simulations to interrogate null expectations for these kinds of experiments. This work has broad implications for infection control strategies and therapeutic techniques that rely on modification of the microbiome, such as probiotics and faecal transplantation.

## Introduction

Human skin is associated with diverse bacterial communities, and this diversity accumulates as a result of repeated invasions and colonizations by microorganisms from exogenous sources (1–5). Touch contact events with surfaces or environmental substrates drive both temporary and longer-term changes in human skin microbial community compositions relative to microbial generation times (6, 7). Empirical studies have characterized skin microbial taxonomic compositions as both variable (6, 8) and stable (9, 10) depending on the time scales under consideration. Yet, the yearly (11–13), monthly (1, 10), weekly (14), and daily (6, 15, 16) intervals that are typically considered by human skin microbiome studies are coarser than those needed to understand transitory changes in the skin microbiome following discrete encounters with environmental microorganisms (17).

Transient dynamics — the dynamical behavior of a system that is different from its long-term, or asymptotic, behavior (18) — are an important aspect of all ecological systems (19, 20), including the human skin microbiome. Far-from-equilibrium dynamics are key to the initial stages of skin microbiome assembly (21, 22) and in the initiation and resolution of dysbiotic events (23–27), such as those that occur during infections or certain disease states (27–29). Understanding and ultimately manipulating the skin microbiome to improve health requires a more complete understanding of the interplay between transient community states reflecting multiple instances of contact and microbial acquisition (6, 10). Empirical characterizations of short-term skin microbial community dynamics following perturbation, that reflect time scales relevant to microbial generation times (30, 31) and human interactions or touch contact events (32, 33), are still needed in attempts to catalogue the ecological and physical processes that influence skin microbial communities.

Here, we report the results of an experiment designed to characterize the short-term dynamics of human skin microbial communities following a single simulated contact event with the microbial communities of (i) farm soil, a relatively high biomass community; and (ii) plant leaves, a lower biomass community. The forearm skin bacterial communities of 16 subjects were sampled prior to a simulated contact event, and again at 2, 4, and 8 hours after contact. In addition, samples were collected from half of the subjects after a wash with soap and water, and from the other half following a 24 hour period with no washing. Results of these experiments identified changes in skin bacterial community taxonomic and phylogenetic compositions indicative of the simulated transfer events, with skin communities more strongly resembling donor communities for a short period before returning to a community composition closer to the pre-perturbed state. The high biomass soil transfer led to detectable changes in community composition that persisted through at least 24 hours in the absence of washing, as well as after washing. In contrast, skin experiencing the lower biomass leaf transfer returned to a composition that was indistinguishable from unmanipulated skin both 24 hours later, and after washing. Simulations of community mixing using sampling models from ecology (34, 35) helped guide conclusions from empirical results, and identified sampling artefacts and unexpected patterns resulting from sampling regimes. Broadly, our results indicate that bacterial abundance in the donor communities is the primary factor determining subsequent detectability and residence times on skin following transfers.

## Methods

The study was performed on 2016 June 20; June 27; and July 13 at the Environmental Studies in Buildings Laboratory in Portland, OR, USA. Sixteen adult subjects between the ages of 18-35 were recruited in Eugene, OR, USA. Eligibility requirements for participation in the study included (i) that the individual was in generally good health, (ii) free of skin conditions or infections, and (iii) had not taken antibiotics within the prior 6 months. Subjects were asked to refrain from bathing or applying topical items to the skin for a 12-hour period preceding the experiment. This study and its associated research protocols were approved by the IRB at the University of Oregon on 2013 December 23. All researchers assigned to this protocol were CITI certified to work with human subjects.

### Experimental design and sample collection

We simulated and standardized the direct transfer of microbes from environmental surfaces and substrates to human recipients using sterile FLOQSwab Specimen Collection swabs (Copan Diagnostics; Murrieta, CA, USA). This was done in order to standardize the bacterial propagule size transferred to each subject from each environmental source. Swabs were also used in the subsequent time-series sampling, as they have been demonstrated to be an effective method for sampling microbial diversity of the skin (1). Initial baseline (we refer to this as time 0) microbial community swab samples were obtained at the inner forearm skin site for each subject (Fig. 1). The inner forearm was selected since this area typically encounters many unique environmental bacterial communities during normal activities. Initial baseline swab samples were also collected for the donor bacterial communities of leaves and soil. Specifically, leaf top surface samples were obtained for three common, indoor plants: *Spathiphyllum wallisii* (Peace Lily), genus *Dieffenbachia* (Mother In-Law’s Tongue), and genus *Calathea* (Prayer Plant). The plants used in this study were purchased at nurseries local to Eugene, OR. Soil samples were aliquoted from farm soil (Mohawk, OR, USA) that was 2 mm sieved and passively air-dried.

**Figure 1:**
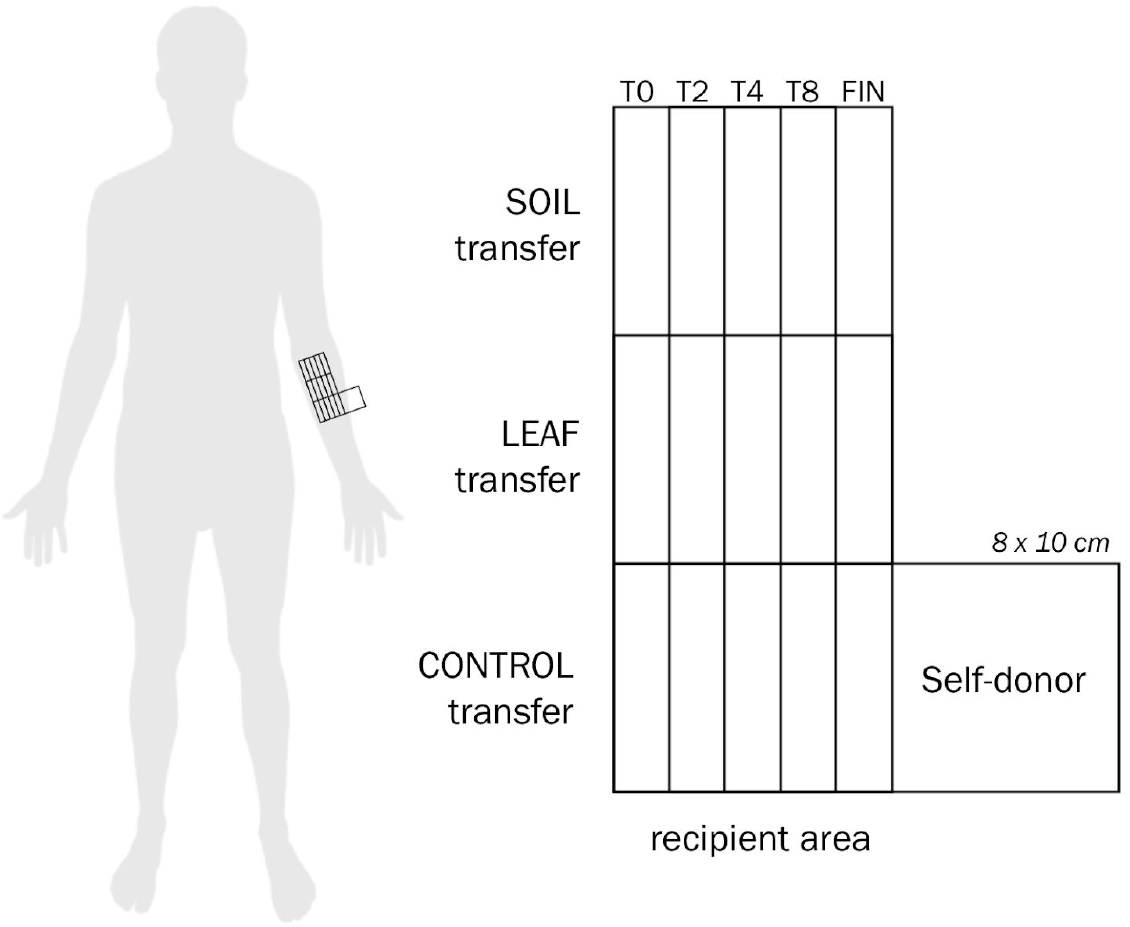
Schematic of sampling area. Sampling areas for each time point (0, 2, 4, and 8 hour, as well as the final 24 hour or wash sample) was randomized between subjects to avoid spatial autocorrelation.

Soil and leaves were selected for several reasons. First, phyllosphere- and rhizosphere-associated microbial communities are sufficiently different from human skin microbiota, so that the coalescence of these communities would generate signals at the level of sequence variants (as opposed to strain-level profiling, that might be needed to characterize human-to-human transfers). Second, these habitats represent a high and low biomass environmental source respectively. By considering these habitats, we were able to simulate the transfer of microorganisms from habitats that were characterized by both more (i.e., soil) and less (i.e., leaves) microbial biomass than human skin, on average (Fig. 2). Finally, people regularly touch dirt and plants, and these microbial sources have been associated with potential human health benefits (36).

**Figure 2:**
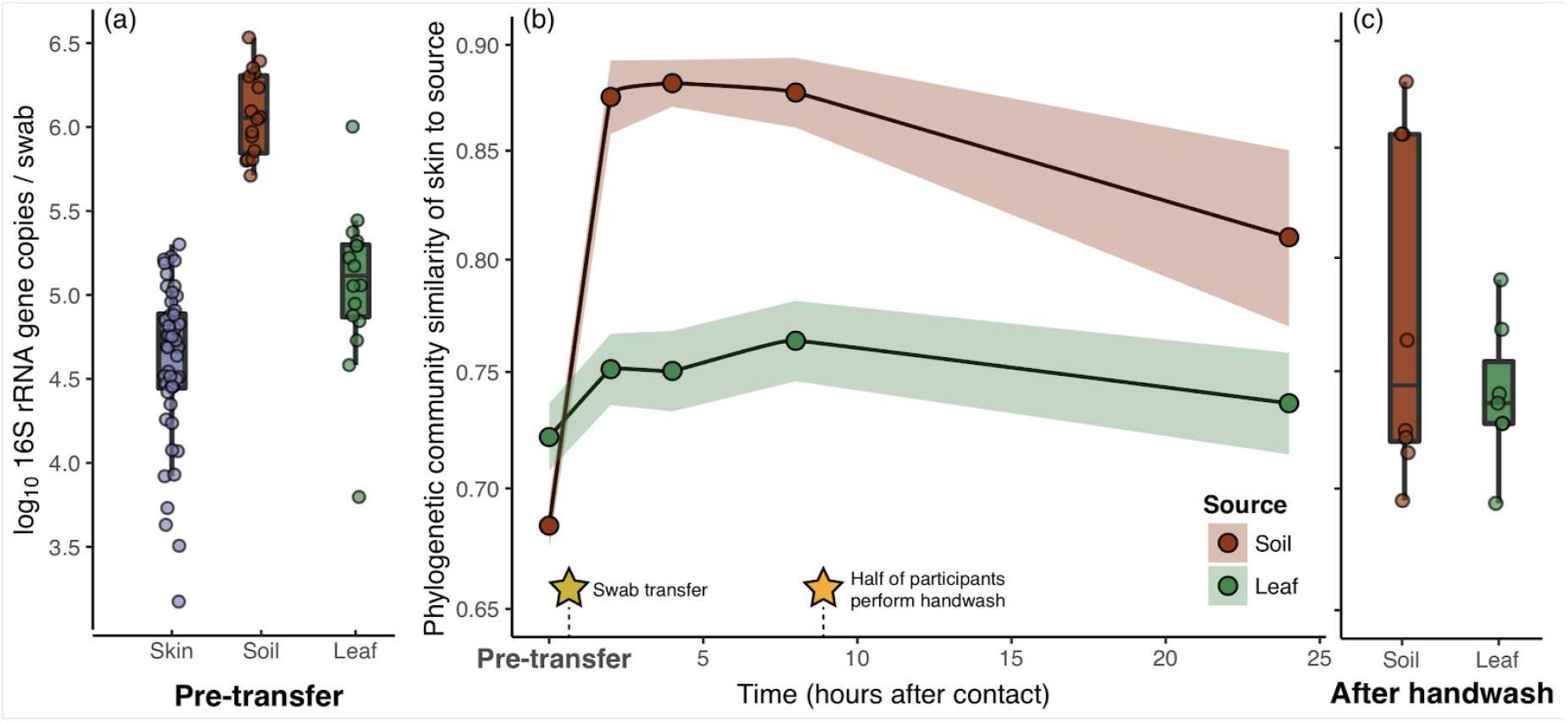
There were significant differences in initial biomass of the two types of donor communities, as well as the initial skin samples before simulated microbial transfer event (a). Phylogenetic community similarities (1 – weighted UniFrac distance) between skin and donor communities after simulated transfer events were elevated for both soil and leaf donor communities, and remained elevated for at least 8 hours; for subjects receiving soil transfers, community similarity remained elevated through 24 hours, though for subjects receiving leaf transfers similarity returned to baseline (b). Community similarity remained elevated for those receiving soil communities after a vigorous soap and water wash, though the subjects receiving leaf communities largely returned to baseline after washing (c).

To standardize the size of the area being sampled and to avoid cross contamination of transfer types, sampling grids were drawn on each subject using ethanol-disinfected custom plastic vinyl stencils and thin-tipped permanent markers to denote “recipient” and “donor” areas of skin for the three transfer types. Five equal and distinct areas were designated for sampling the skin at five time points: before transfer (0); 2-, 4-, and 8-hours post-transfer; and a spot for either a 24-hours post-transfer or post-wash (FIN) sampling time point (Fig. 1). To avoid spatial autocorrelation, the location of the sampling area for each time point was randomized within the sampling grid. After the baseline samples of donor and recipient communities were collected, microbial communities were immediately transferred from a donor plant leaf and from an aliquot of farm soil to the dry inner forearm of an individual human recipient subject. Skin-to-skin control transfers were also performed using a designated area of dry skin on the arm to inoculate an adjacent area of dry skin.

To perform each transfer, a sterile swab was first dipped into a sterile, saline solution (0.15 M NaCl; 0.1% Tween20) and then rid of excess moisture by flicking the swab carefully. For transfer of the plant donor community, the swab was then applied to a plant leaf across a 10×10 cm area and rotated while swabbing firmly for approximately 10-15 seconds. The swab was then applied to the recipient area of skin and again rotated while swabbing firmly for approximately 10-15 seconds. A different donor leaf was selected and swabbed for each human subject volunteer. Subjects 01-06 received a leaf donor community from *Spathiphyllum wallisii*, Subjects S07-S12 received a leaf donor community from *Dieffenbachia*, and Subjects S13-S16 received a leaf donor community from *Calathea*. Plant identity was, however, non-significant in all downstream analyses. For transfer of the soil donor community, the protocol was identical except that after the swab was dampened it was briefly dipped into a ∼20 mL aliquot of farm soil to collect soil. The swab was then applied firmly and rotated for 10-15 seconds on the recipient skin area.

After the transfer procedure was complete a sterile gauze dressing was lightly placed and taped to cover the area with minimal occlusion and replaced between sampling periods. Subjects remained sedentary in a climate-controlled chamber for 8 hours. For each of the 3 transfer types on each subject at the requisite time point 2, 4, 8, and 24-hours post-transfer (and post-wash), a non-destructive swab sample was collected using a sterile swab (dampened with sterile saline solution: 0.15 M NaCl; 0.1% Tween20) applied to the skin and rotated while swabbing firmly for approximately 10-15 seconds. All baseline transfer swabs (T0) and following time-series swab samples (T2, T4, T8) were collected and frozen at −20C for subsequent DNA extraction.

### Skin washing and 24 hour treatment

To achieve both a post-wash (TW) sampling time point and a 24-hour time point (T24), we asked half the subjects to wash the recipient skin area with Castile soap and gently pat dry with sterile paper towels just after the 8-hour sampling time point for immediate re-sampling. The remaining eight subjects did not wash the recipient skin area but were sampled the following day for a 24-hour post-transfer sampling time point. Between the time that the subjects left the research facility and returned the following day for a 24-hour sampling time point, they were asked to refrain from thoroughly wetting the now-exposed inner forearm area, but otherwise were asked to resume daily activities. TW and T24 swab samples were also collected and frozen at −20C for subsequent DNA extraction.

### 16S rRNA gene sequencing

The DNA from all samples was manually extracted using the MoBio PowerLyzer PowerSoil DNA Isolation Kit according to manufacturer’s instructions. Amplicons of the V3-V4 region (319-806) of the 16S rRNA gene were prepared in 50 µL PCR reactions with one PCR step using dual-barcoded primers (see description in data deposition (37)), cleaned with Ampure beads, quantified using Quant-iT dsDNA assay kit, and pooled with equal concentrations of amplicons using an Eppendorf epMotion 5075 robot. Libraries were sequenced on an Illumina MiSeq generating 250 bp paired end reads.

Illumina sequence data were filtered, trimmed, and denoised using the DADA2 v1.5.2 statistical inference algorithm (38, 39), which identifies ribosomal sequence variants (RSVs). Forward reads were trimmed and truncated at 10 nt and 240 nt, and each read was required to have fewer than three expected errors based on quality scores. Taxonomy was assigned to RSVs using the RDP classifier implemented in DADA2 and the Silva version 123 reference database (40), with an 80% bootstrapped threshold for retaining classifications. We omitted sequence variants classified as chloroplasts or mitochondria, and those that were unclassified beyond the kingdom level from analyses. Putative contaminants were removed following the suggestion of Nguyen et al, (41) by subtracting the number of sequences of RSVs present in negative PCR and extraction kit controls from sequence counts in experimental samples.

### Estimating microbial counts in source communities

We estimated the size of each transfer propagule as total counts of 16S rRNA gene copies per swab (a proxy for absolute bacterial abundances) for soil and leaf source communities using real-time quantitative PCR (qPCR; Applied Biosystems StepOnePlus System). We estimated total counts of 16S rRNA gene copies per mg of soil (approximately 0.0075 g of soil from the original swab transfer) and per area of leaf swabbed (*ca.* 10 × 10 cm area of the top of a single leaf of the donor plant). The reaction mixture was prepared according to guidelines provided by ABS PowerUp SYBR Green PCR Master Mix for a 20 µL reaction and ran in triplicate: ABS PowerUp SYBR Green PCR Master Mix (10 µL), 10 µM Total Bacteria F SYBR Primer 5’-gtgStgcaYggYtgtcgtca-3’ (0.8 µL), 10 µM Total Bacteria R SYBR Primer 5’-acgtcRtccMcaccttcctc-3’ (0.8 µL), PCR grade water (6.4 µL) and 2 µL of undiluted DNA template (42). We also used the suggested ABS PowerUP thermocycling conditions for primers with T_m_ < 60°C: initial denaturation for 2 min at 50 °C, 2 min at 95 °C; 40 cycles of 15 sec at 95 °C, 15 sec at 60 °C, 60 sec at 72 °C; followed by a melt curve in the range of 60 °C to 95 °C. Standard curves were generated using 10-fold serial dilutions of synthetic 167 bp gBlocks Gene Fragments (Integrated DNA Technologies, Coralville, Iowa, USA) of the same region amplified with the above primers, with known gene sequence copy numbers. To correct for differing reaction efficiencies across the different source community sample types (43) we used LinRegPCR (44) to quantify 16S rRNA gene copies per swab from fluorescences values.

### Statistical analysis

All analyses were conducted with the statistical programming language, R (45). Differences between qPCR-based estimates of initial taxon abundances were examined using analysis of variance, ANOVA, and Tukey’s post-hoc test.

We created a phylogenetic 16S gene tree for all RSVs in our data using a maximum likelihood GTR+Gamma phylogenetic model in FastTree (46), following Callahan et al. (38). This 16S gene tree was used to calculate weighted Unifrac distances, which measure phylogenetic and compositional dissimilarity between pairs of communities (47). To examine the change in *β*-diversity between donor communities and re-sampled skin communities through time, we fitted generalized linear mixed effects models with beta error and a logit link function (48), to estimate the rate of change in Unifrac distances between the donor and the resulting mixed communities through time, while accounting for the random effect of individual subject. Beta mixed models were fitted using the *glmmTMB* library in R (49).

Generalized linear mixed effects models with binomial error and a logit link function (50) were used to estimate changes in the probability that an RSV persisted after transfer, and its dependence on initial taxon absolute abundance in the donor community. Microbial persistence was scored as a binary measure, with persistence being operationally defined as the detection of a taxon during a census and all prior censuses (i.e., if a taxon was detected at 2, and 4 hours after transfer, it is said to have *persisted* until the four-hour census). Using the *lme4* library in the R statistical programming environment, our model fits to the binary persistence variable as a function of the mean calculated initial 16S gene copy number for that taxon in the donor community (i.e., estimates of absolute abundance from qPCR assays), the time of sampling, and the interaction between the two, with subject included as a random effect.

We examined phylogenetic structure within each community for each subject. By utilizing the standardized effect size (SES) from a permutational test of the Mean Nearest Taxon Distance (MNTD) (51) as implemented in the *picante* package in R (52). Briefly, the unweighted 16S gene tree created above was used to calculate the mean distance within the tree to the nearest taxon of a shared classification (in this case, presence on the skin after time *t*); this is compared to the distribution of MNTDs resulting from 10^3^ permutations of the same number of randomly drawn tip labels across the same tree, and the SES is calculated as the difference between the observed MNTD and the permutational mean MNTD, over the standard deviation of the permutational MNTDs. The *P*-value is calculated as the proportion of permutations in which the observed MNTD is less than that of the randomly selected tips. If there is no phylogenetic structure in the grouping, we expect that the MNTD should be, on average, no different from the MNTD of a random draw of the same number of tips. To test assumptions of a phylogenetic signal of abundance, we subsetted taxa into abundance classes (the top 1%, 5%, 10% and 25% of taxa), and tested the MNTD SES for these groupings, as well.

To elucidate particular groups of related organisms and specific taxa that discriminate skin communities before and after experimental transfers, we used phylogenetic tree-based sparse linear discriminant analysis (sLDA) as a feature selection tool, to identify whether individual taxa or groups of related taxa discriminated across our temporal sampling. This analysis is described by Fukuyama et al. (53). Briefly, using the same 16S gene tree above, we generated two sets of features, one made up of the relative abundances of all taxa present as tips on the tree, and the other including all nodes within the tree; for the set of nodes, feature values were the sum of the relative abundances of all taxon tips descended from that node. These feature sets were centered, and used as the input to the implementation of sLDA in the *sparseLDA* package. The optimal number of model predictors and sparsity parameter were determined by 3 repeats of 5-fold cross-validation.

### Sampling simulation modeling

Building on previous sampling models (34, 35), we developed null expectations for changes in taxon abundances and community composition following the experimental mixing and resampling of bacterial communities (54, 55). This allows us to predict qualitative differences in perceived changes in taxon abundance and β-diversity when sampling from mixed communities as compared to samples from the two communities before mixing. Specifically, the model predicts changes in the detection rates, and therefore the apparent abundances, of taxa in identical communities with and without the addition of new taxa through the transplantation of a fixed portion of an independent community. The model is also applied to predict changes in observed β-diversity between such identical communities before and after mixing with an independent SAD as it varies with sampling depth. The addition of large number of new taxa, some of which are numerically abundant, imposes certain constraints upon the sampling of these mixed abundance distributions. Here, we seek to investigate these constraints using a permutational simulation modeling approach, generating null expectations regarding those constraints.

Following previous work (35), our model begins with assuming two independent underlying log-series sequence abundance distributions (SADs) describing the abundances of 16S rRNA gene copies originating from S bacterial taxa in a community. We assume no members in common between the two communities, and log-series distributions for SADs (56), with probability parameterized for each iteration to be between the observed probabilities for the skin and soil SADs in this experiment.

To study perceived changes in relative abundance in rare taxa, we assumed 50% of individuals in donor communities were mixed with 100% of individuals from recipient communities. This new mixed SAD and the initial recipient SAD were then sampled at a fixed percentage (half of individuals present), and converted to relative abundances in the sample, to mirror the type of observations possible in metabarcode amplicon studies. Apparent log_10_-fold change in relative abundance from initial sample to final sample was calculated for all simulated taxa sampled in both communities. This variable is a measure of change in the relative abundance of target taxa after mixing and re-sampling, and is often conflated with change in the true abundance of taxa in time series sampling. We performed 10^4^ iterations of this sampling procedure, independently drawing the probability parameter from a uniform distribution bounded by experimental outcomes (see Results). To more closely simulate metabarcode sequencing, simulations were also run using fixed number sampling (10^3^) rather than fixed percentage sampling, which yielded qualitatively similar results (Supp. Fig. 1a). Additionally, a third set of simulations was performed using the recipient community as a self donor, to provide a similar control to the experiment described above (Supp. Fig. 1b).

To evaluate effects of sampling depth on perceived β-diversity between initial and final communities, donor and recipient SADs were generated as above, and a range of sampling depths were generated for each simulation. For each sampling depth within each simulation, β-diversity was calculated as the Bray-Curtis distance between the initial recipient and final mixed communities. We performed 10^4^ iterations of this sampling procedure, with 20 sampling depths per simulation, independently drawing the probability parameter from uniform distributions bounded by experimental outcomes, as above, and sampling depth from a uniform distribution of depths bounded on the upper limit by one quarter of the resulting individuals in the SAD. We summarized these results using plots of log_2_-transformed ratios of observed to true β-diversities as a function of sampling depth. Each point on these plots represents the results of a single simulation at a single sampling depth, and positive values on the y-axis of these plots indicate that observed β-diversity was inflated relative to the true β-diversity.

## Results

Sampled bacterial biomass densities (bacterial 16S rRNA gene copies per swab) differed between human skin, soil, and leaf communities (Fig. 2a), prior to simulated microbial transfers (ANOVA of log_10_-transformed 16S rRNA gene counts; *F_2,77_* = 68.64, *P* < 0.001). Soil communities were characterized by *ca.* an order of magnitude more 16S rRNA gene copies per swab compared to skin communities (ANOVA; *β_soil_* = 1.48, *P* < 0.001), whereas the difference between skin and leaf communities was comparatively smaller (ANOVA; *β_leaf_* = 0.47, *P* < 0.001).

### Beta-diversity dynamics depend on donor community

#### Soil

Samples from participants who received soil bacterial communities during simulated transfers (i.e., high biomass transfers) strongly resembled the donor community compared to the pre-transfer state, even 24 hours post transfer (Fig. 2b; Table 1). Communities observed on the skin remained significantly more similar to donor communities, even after washing (Fig. 2c; ANOVA *F_1,29_* = 22.48, *P* < 0.001).

**Table 1:**
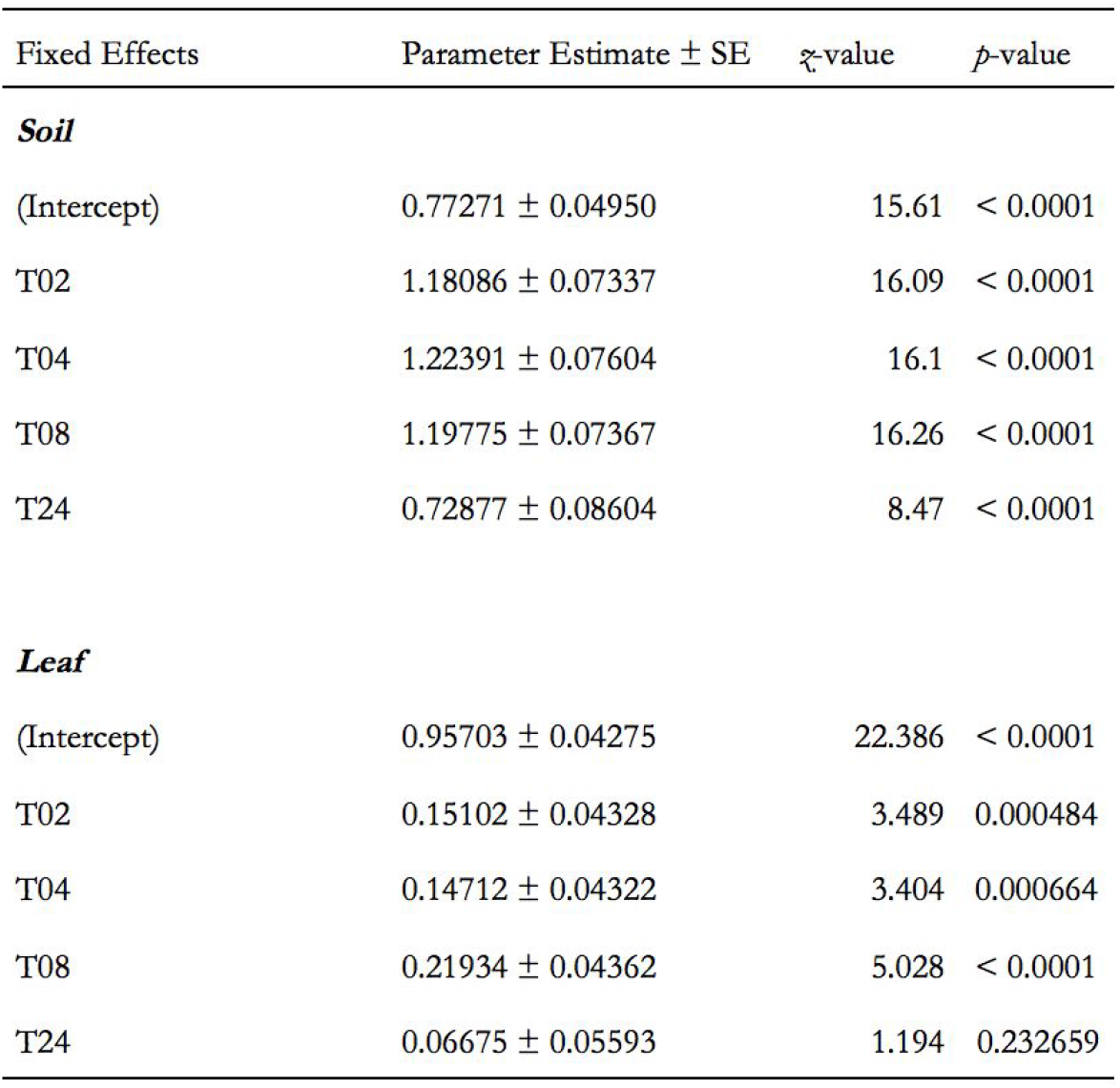
Results of generalized mixed-effect models with beta error (glmmTMB) to test the effect of time since simulated microbial transfer event on the Unifrac distance between skin communities. The model was of the form **Y ∼ *B_1_*(time) + *G* + *E***, where Y is Unifrac distance, *B_1_* is a regression coefficient, *G* is a random effect of subject to account for host-specific variation, and *E* is a vector of errors.

#### Leaf

Samples from participants who received leaf microbial communities (i.e., low biomass transfers) were also significantly more similar to the donor community compared to baseline, although this change was commensurately smaller compared to the soil transfer experiments. Skin communities receiving leaf-associated bacteria returned to baseline by the 24 hour census (Fig. 2b; Table 1). Leaf transfer communities also returned to baseline after washing, although there was marginal increased similarity relative to baseline after washing (Fig. 2c; ANOVA *F_1,29_* = 4.666, *P* = 0.040).

### Biomass drives response

#### Soil

For the comparatively high-biomass soil community, we found a significant effect of initial biomass (Fig. 3a). Specifically, we observed an impact of initial 16S gene copy number on whether or not donor taxa were detected after the simulated transfer event, or persisted (were re-detected) through time (Table 2). For the purposes of this study, we define *persistence* as a microbe being detected at a time point (n), and also having been detected at the previous time point (n-1). A similar effect on the initial skin community was also detected: there is a significant effect of initial 16S gene copy number on whether or not skin taxa were re-sampled after the simulated transfer event (Table 2). Unexpectedly, we also saw an apparent increase in the relative abundance of rare skin-associated taxa re-sampled after simulated microbial transfer events (Fig. 3c).

**Figure 3:**
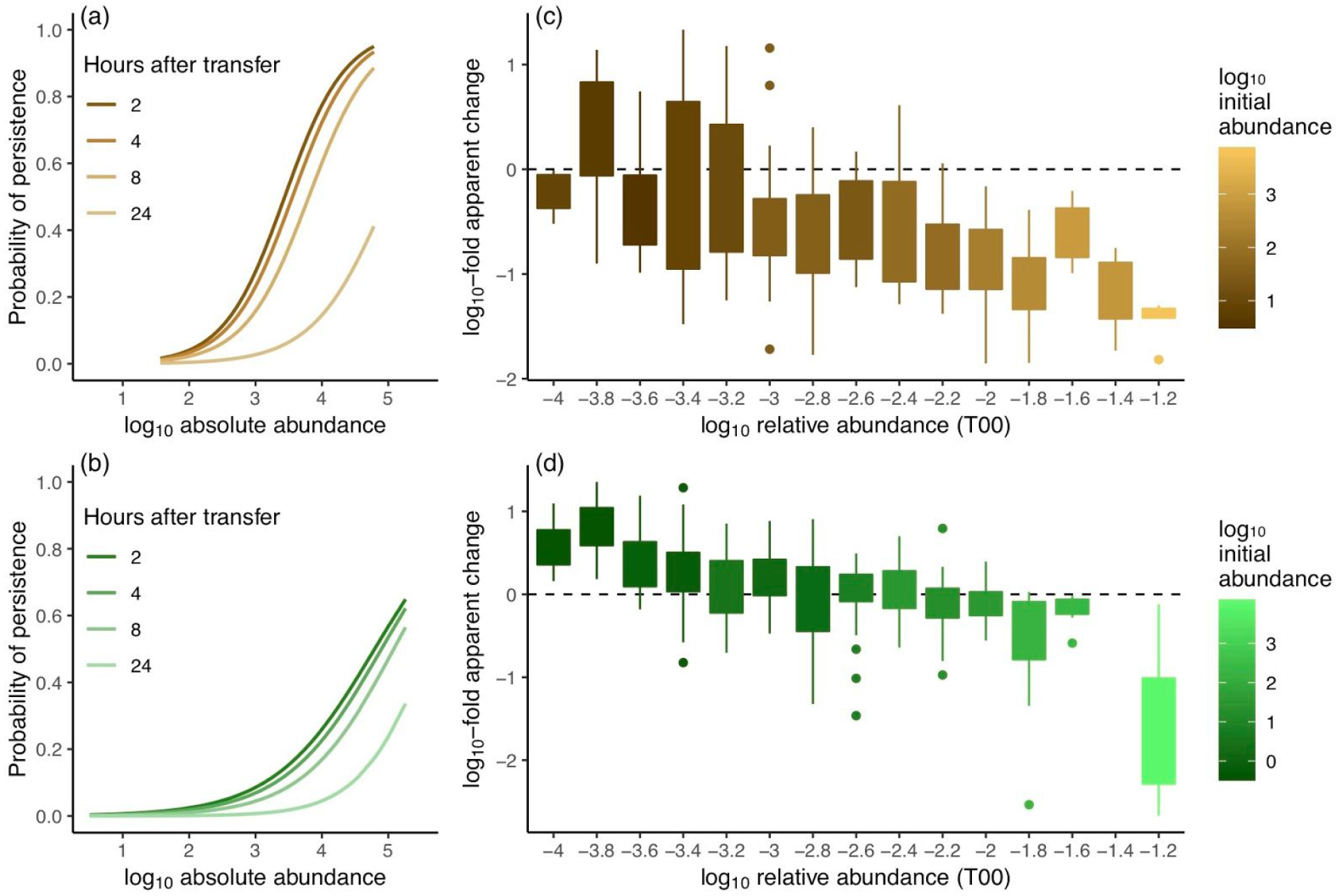
Initial *absolute abundance* in the donor community (as estimated by 16S copy number measured by qPCR) and *time* both have predictable effects on the probability of persistence (i.e., the probability of detection at sequential time points on the skin) in both soil (a) and leaf (b) simulated microbial transfer events. Probability of persistence for both soil (a) and leaf (b) transfers are estimated from GLMM model fits (Table 2). For skin-associated microbes sampled before simulated microbial transfer event (at time T00), there is a consistent pattern of apparent increases in relative abundance for rare taxa when they are re-sampled two hours after transfer (at time T02) for both the comparatively high-biomass soil (c) and the low-biomass leaf (d) simulated microbial transfer event.

**Table 2:** Results of generalized mixed-effect models with binomial error (lme4) to test the effect of log_2_-transformed absolute abundance and time since simulated microbial transfer event on the persistence (i.e., sequential detection) of microbes on the skin. The model was of the form **Y ∼ *B_1_(A)* + *B_2_(T)* + *B_3_(A*T)* + *G* + *E***, where Y is the binomial probability of re-detection on the skin; A is log_2_-transformed initial absolute abundance; T is time; *B_1_*, *B_2_*, and *B_3_* are regression coefficients; *G* is a random effect of subject to account for host-specific variation; and *E* is a vector of errors. Scaled and centered values were used for both fixed effects.

| Fixed Effects | Parameter Estimate ± SE | $z$-value | $p$-value |
| --- | --- | --- | --- |
| ***Soil*** |  |  |  |
| $\log_2$ (abundance) | 1.010 ± 0.02161 | 46.72 | < 0.0001 |
| time | -1.046 ± 0.02332 | -44.87 | < 0.0001 |
| $\log_2$ (abundance) * time | -0.075 ± 0.02278 | -3.29 | 0.001 |
| ***Leaf*** |  |  |  |
| $\log_2$ (abundance) | 0.840 ± 0.05717 | 14.70 | < 0.0001 |
| time | -1.189 ± 0.14605 | -8.14 | < 0.0001 |
| $\log_2$ (abundance) * time | 0.123 ± 0.06608 | 1.86 | 0.063 |
| ***Skin (with Soil transfer)*** |  |  |  |
| $\log_2$ (abundance) | 1.047 ± 0.05030 | 20.806 | < 0.0001 |
| time | -0.948 ± 0.09272 | -10.226 | < 0.0001 |
| $\log_2$ (abundance) * time | 0.013 ± 0.05480 | 0.245 | 0.806 |
| ***Skin (with Leaf transfer)*** |  |  |  |
| $\log_2$ (abundance) | 1.515 ± 0.06281 | 24.089 | < 0.0001 |
| time | -1.123 ± 0.09175 | -12.237 | < 0.0001 |
| $\log_2$ (abundance) * time | -0.049 ± 0.06554 | 0.744 | 0.457 |

#### Leaf

The effect of initial biomass in the donor community was also observed in the lower biomass leaf community transfer (Fig. 3b; Table 2), though the effect was weaker than for the soil, likely due to the comparatively smaller initial total biomass and richness in the leaf donor pool. The effect of initial 16S gene copy number on whether or not skin taxa were re-sampled after the simulated transfer event was stronger with transfer from the lower-biomass leaf community than from the soil community (Table 2). The transfers from leaves also led to apparent increases in relative abundance of skin taxa that were re-sampled after the simulated transfer event (Fig. 3d).

### Phylogenetic considerations

#### Soil

Interacting with these effects of biomass, microbes from the soil donor communities re-sampled on the skin after simulated microbial transfers were more phylogenetically clustered—i.e., had reduced *mean nearest taxon distances* (51)—than expected by chance (2-hour post-transfer: MNTD on 999 permutations, by subject; *z-*bar = −2.343, *P*-bar = 0.010). In fact, all time points sampled were significantly clustered, save the initial donor community and the soil microbes recovered after washing (Supp. Fig. 2a). However, there is a strong phylogenetic signal of initial abundance in the donor soil communities (top 10% of taxa, MNTD on 999 permutations, by sample; *z*-bar = −4.447, *P*-bar < 0.001). As more abundant taxa are more likely to be detected (and re-detected) on the skin at subsequent time points (Fig. 3), and phylogenetic clustering correlates more strongly with abundance, it is likely that the resulting clustering is due to selection through the transfer process for more abundant taxa (Supp. Fig. 3).

Additionally, microbes found on the skin before simulated microbial transfer that were re-sampled after the transfer event were marginally more phylogenetically clustered than expected by chance for (2-hour post-transfer, soil: MNTD on 999 permutations; *z* = −1.512, *P* = 0.029). As with the donor communities, the initial skin microbial communities also displayed strong phylogenetic clustering with abundance (top 10% of taxa, MNTD on 999 permutations, by sample; *z*-bar = −1.635, *P*-bar = 0.003), so that the marginal effect of simulated microbial transfer from soil is easily explained by the selection for abundant skin-associated taxa imposed by sampling from the mixed the community with fixed sampling effort relative to the initial baseline sampling events.

Linear discriminant analysis with sparsity constraint (sLDA) on the matrix of tips and nodes from the microbial community 16S trees was used to select clades or single taxa responsible for observed β-diversity changes between initial skin sampling and the first post-transfer sample (53). This method identified 3 clades comprised of a total of 216 RSVs in soil transfers that strongly discriminated communities across sampling time points based on their feature loadings on the discriminating axis (Supp. Fig. 4a).

#### Leaf

Microbes from the commensurately lower biomass leaf donor communities re-sampled on the skin after simulated microbial transfers were *not* significantly phylogenetically clustered at any time point (Supp. Fig. 2b), despite the persistence of a strong phylogenetic signal of initial abundance in the donor leaf communities (top 10% of taxa, MNTD on 999 permutations, by sample; *z*-bar = −2.983, *P*-bar < 0.001).

Additionally, microbes found on the skin before simulated microbial transfers that were re-sampled after the transfer event were no more phylogenetically clustered than expected by chance for (2-hour post-transfer, leaf: MNTD on 999 permutations; *z* = −1.107, *P* = 0.132). The lower biomass present in the leaf simulated microbial transfer appears not to have been sufficient to bias the re-sampling towards microbes with high initial abundance enough to create significant clustering in the downstream samples.

The linear discriminant analysis (sLDA) method also identified 3 small clades comprising a total of 44 RSVs in leaf transfers that strongly discriminated communities across sampling time points (Supp. Fig. 4b).

## Discussion

Here, we empirically characterized fine-scale dynamics after a whole-community microbial transmission event to human skin, describing the relationship between initial abundance in the donor community and persistence through time. While it has been demonstrated that such contact events lead to an increase in skin microbial diversity (57), and that the microbiome of the skin is both highly diverse and highly variable across longer time scales (6), the fine-scale dynamics of microbial transmission through such contact events has not been previously characterized.

We expected that simulated microbial transfers would move a representative sample of the donor community to the recipient: that is, if the movement of microbes is determined by density-independent *per capita* processes, we would expect that microbes that are more abundant in the donor community would be transferred more frequently, and that the identity or phylogeny of microbes in the donor community would be unrelated to their probability of being transfered (beyond the way that identity or phylogeny is related to abundance in the source community). We would also expect that effects on the recipient skin community would reflect sampling from a larger, more diverse species pool, leading to apparent loss of rare taxa from the pre-transfer sampling. This is, indeed, largely what we found, although with some important caveats.

### Biomass and Persistence

Initial biomass in the donor community drives trends in detection of exogenous microorganisms at later sampling times (Fig. 3a–b). Additionally, we found that the probability of persistence on the skin (i.e., the probability of being detected at subsequent, sequential time points) scaled predictably with time since simulated transfer event: for every 14.5 hours after simulated microbial transfer event from soil, an additional order of magnitude of microbial abundance in the donor community is required to maintain equal probability of persistence. This effect was qualitatively the same for the leaf transfer events, with 22.5 hours post-transfer requiring an additional log-fold increase in microbial abundance to maintain constant probability of persistence. Despite the vast differences in initial community membership and biomass, these estimates are within an order-of-magnitude; it will be necessary to determine if this relationship holds across a broader range of subjects and donor communities, but if this decay relationship is at all generalizable, it suggests an initial basis for modeling interacting transient assemblages of exogenous microbes on human skin. This has important implications for hospital infection control strategies (58, 59), bacterial disease transmission (33), and developing strategies to make use of microbial mediated immune system modulation (60–62).

Most microorganisms remained on human skin for only several hours, whereas a small number of numerically abundant and phylogenetically related taxa were detected on subjects for at least 24 hours, and after a thorough washing with soap and water. The probability of successful colonization by microorganisms from environmental sources depends largely on their absolute abundances in source habitats (63–65). Our results characterize the transient timescales of exogenous microbial acquisition by human skin, and reveal that the probability of persisting through time depends on initial biomass in the source communities. This drives changing β-diversity between skin microbial communities after the transfer event and the initial donor microbial communities. The practical effect of this is that the propagule pressure for abundant taxa emigrating to the skin is higher; on longer time scales, this implies that the probability of colonization (as distinct from the probability of *acquisition*, studied here) is likely to be higher for those taxa that maintain high abundance in source communities that are often contacted (66–68).

### Community Coalescence and Non-equilibrium Dynamics

Touch contact with substrates such as soil or plants, which bear their own complex assemblages of microbial residents, may lead to the wholesale transmission of microbial communities, a phenomenon known as *community coalescence* (69). Microbial community coalescence theory seeks to fill a gap in ecological theory developed for macroscopic eukaryotic organisms: namely, that it is only possible to have wholesale community-community interactions when those communities are microbial (70). While it has been demonstrated that such community interactions can lead to co-recruiting of taxa through community co-selection mechanisms (through bottom-up interactions, where the presence of certain facilitator community members help to recruit dominant members of a transfered community after a coalescence event; or top-down interactions, where the transfer of dominant members of a community leads to co-selection of community members that depend upon that dominant member) (71), it is unclear if such dynamics are present on the human skin after a whole-community transmission event. Our observations of transient time scales after a community coalescence event would seem to indicate that physical processes are the primary drivers of microbial dynamics after contact events.

It is possible that regular acquisition through dispersion (i.e., contact) events and the transient dynamics of resetting the microbiome are enough to explain the increased variation observed on the skin relative to the gut and oral microbiomes (6). That is, the skin microbiome may be *always* in a non-equilibrium state, ‘recovering’ from a recent coalescent event. Differential persistence through time for different microbes (potentially driven by differential abundance in the source communities) means that there will be complex behavior resulting from the interaction of different time scales: the time scales on which microbes are lost (which vary by microbe, and by other disturbance events like washing), and the time scales on which new contact events happen. This sets up a situation in which the only way to explain the microbial community present on the skin at a given moment is not through asymptotic or stable equilibria (72), but through transient dynamics of multiple, interacting coalescence events — the current microbial assemblage as the result of the combination of multiple contact events across time, with each microbe acquired having its own particular decay curve, all of which must be considered together to explain any given assemblage. These results have implications for the study of hand hygiene (73), given that biomass in the source community moderated the ability of skin communities to return to baseline after washing events.

### Sampling Effects

There is, however, a danger of assuming that the patterns observed in microbiome coalescence experiments such as this one are ecological in nature. Here, we used sampling simulation models (see ‘Methods: Sampling simulation modeling’, above) to define null expectations (54) for meta-barcode sampling from combined species abundance distributions. Briefly, we combined independent simulated sequence abundance distributions, and then re-sampled from those combined distributions, comparing samples to the initial distributions; this mirrors the experiment described herein. For each of the 10^4^ iterations of these simulations, we measured the apparent change in relative abundance (that is, the difference between the relative abundance of simulated taxa from the final mixed sample and from the initial sample), and the β-diversity at multiple sampling depths. To visualize null expectations around changes in relative abundance, we plotted the log_10_-fold apparent change as a function of the relative abundance in the initial sample (Fig. 4b). To visualize null expectations around β-diversity measurements for mixed communities with varying sampling depth, we plotted the log_2_-transformed ratio of the observed Bray-Curtis distance (calculated from the sample) to the true Bray-Curtis distance (calculated from the full simulated data) as a function of the sampling depth (Fig. 4a).

**Figure 4:**
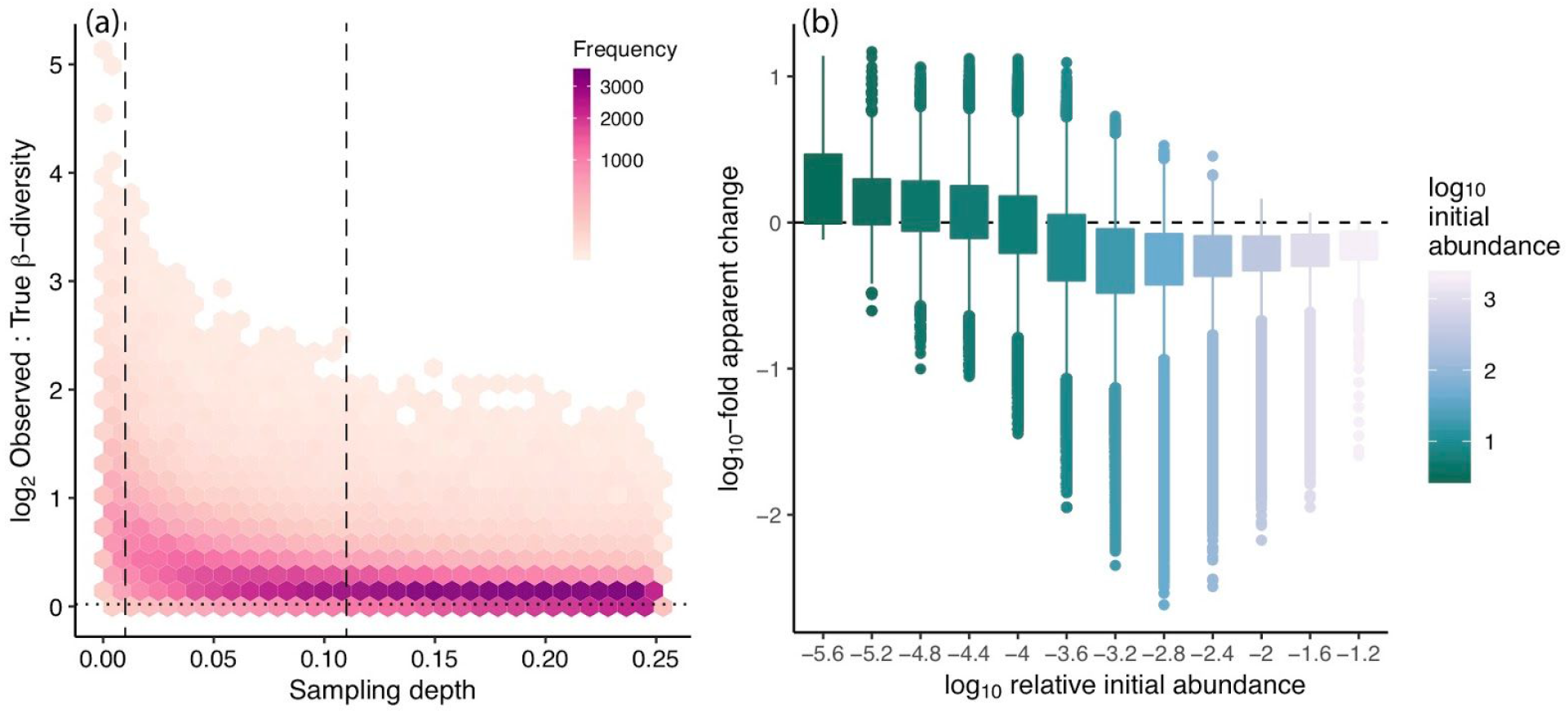
Sampling simulation models (10,000 permutations) indicate that estimated β-diversity from mixed communities will likely be elevated relative to true β-diversity when sampling depth is less than 5% for both samples (a). Vertical dashed lines represent 1% and 11% sampling, reflecting measured sampling depths in this study for soil and leaf microbial communities, respectively. The average sampling depth for the low-biomass initial skin community samples was 43% (not shown). Similarly, comparisons between initial samples and samples from mixed communities indicate that an apparent increase in abundance of rare taxa is an expected, though artefactual, effect of resampling from mixed communities (b; see also Supp. Fig. 1).

Our models provide some null expectations that may seem counterintuitive, including inflated β-diversity estimates if sampling depth is low (Fig. 4a) and an apparent increase in abundance of resident members of the rare microbiome (Fig. 4b), which qualitatively matches our empirical observation (Fig. 3c–d). The understanding that such apparent changes in relative abundance is not anomalous, but expected, is particularly important given the admonition that the members of the rare biosphere often “display specific and sometimes unique ecology” (Lynch and Neufeld 2015). Without an understanding of the particular constraints that resampling from mixed species pools places upon patterns of observed relative abundance distributions, it would be easy to interpret these observations as representing ecological differentiation in the rare biosphere.

We also observed apparent phylogenetic signal in the microorganisms transferred from the soil to the skin (Supp. Fig. 2), but this effect appears to be a reflection of the relationship between abundance in the donor community and probability of transfer and persistence (Fig. 3). Other studies have reported phylogenetic clustering of microbial communities in various situations (for example: in soils of various successional ages (74), in shrimp of differing disease states (75), and in rhizosphere microbiomes (76)); indeed, we find that the soil, leaf, and skin communities are clustered relative to the tree of all taxa (data not shown). Our results highlight that such analysis must be undertaken carefully — while we *do* find evidence that transferred microbes are more phylogenetically clustered than expected by chance, which could be interpreted as evidence that there is ecological differentiation across the microbial tree of life with regards to their ability to be acquired by or persist upon human skin, the effect is more parsimoniously explained by examining the way that biomass is related to phylogenetic clustering. As with any ecological discipline, the study of the ecology of the human microbiome and coalescence as it might relate to human-associated microbial communities must be careful to disentangle sampling artefacts from true ecological explanations for observed trends (35, 55, 77, 78).

### Skin Microbiome Dynamics and Disease

It is important to understand the role of transient dynamics in human skin microbial assemblages because there is mounting evidence that diverse microbial exposures may play key roles in reducing the risk of immune-mediated diseases (79). The so-called ‘Hygiene Hypothesis’ postulates that a lack of exposures to microorganisms, particularly early in development (80), increases susceptibility to allergic diseases by suppressing the natural development of the immune system (81, 82). Humans in industrialized nations spend upwards of 90% of their time indoors (83), which significantly moderates their microbial exposures. Indoors environments tends to include more human-associated taxa than outdoor environments (84–86), and tend to have reduced biomass as compared to soil or plant-associated communities (15, 87). This will mean that while indoor exposures may display reduced residence times on the skin for incoming environmental microbes (through reduced inoculating biomass; Fig. 3), it is probable that they will also display increased probability of long-term colonization events or other human-microbe interactions (15). Because of constant microbial shedding by building occupants (88), it is probable that indoor exposures will display increased similarity with the existing skin microbiome, which may lead to reduced divergence with indoor-sourced microbial coalescence events. One testable hypothesis that results from this line of reasoning is: are the skin microbiomes of people from industrialized nations, or people that lead largely indoor lives, more stable?

Furthermore, there are numerous therapies that rely on direct modification of the human microbiome (usually, the gut microbiome) through intentional coalescence events, for example faecal microbiota transplants (FMT) (89), and probiotic therapies (90). The use of probiotics for digestive health similarly involves the melding of species abundance distributions and has been widely applied, despite the fact that the transient dynamics (90) and method of action (91–93) of probiotics application are still poorly understood. One community of microorganisms is added to another, and the resulting community displays changes in structure and function as a result. The colonization of existing habitats due to the application of probiotics has been documented over short time scales (90), but the ecology and long-term dynamics are not well characterized (93). An application of ecological theories for transient dynamics (72) and coalescence events (69) may aid in the framing of new experimental questions examining the method of action of probiotic therapies.

## Acknowledgements

This work was funded by a grant to the Biology and the Built Environment Center from the Alfred P. Sloan Foundation Microbiology for the Built Environment Program (no. G-2015-14023). The funders had no role in study design, data collection and analysis, decision to publish, or preparation of the manuscript.

The authors would like to thank Dr. Jessica L. Green, who provided laboratory space, mentorship, and aid in study design while serving as a dissertation co-advisor (with BB) for AB, as well as Dr. Roxana J. Hickey, who provided invaluable help with design and execution of the experiment. We would also like to thank Hannah Wilson and Clarisse Betancourt for sample processing and sequence library preparation, and Maggie Weitzman (University of Oregon’s Genomics & Cell Characterization Core Facility) for sequencing and advice. A special thanks to the Energy Studies in Buildings Laboratory (ESBL) for use of their climate-controlled chamber, and to Jason Stenson in particular for his assistance in using the chamber. We also thank Marshall Bockrath for statistical consulting, and Isabelle O’Neal for graphic design assistance.

## Author Contributions

RV analyzed the data, wrote the paper, prepared figures and/or tables, reviewed drafts of the paper; AKF analyzed the data, wrote the paper, prepared figures and/or tables, reviewed drafts of the paper; MM conceived and designed the experiments, reviewed drafts of the paper; ACB conceived and designed the experiments, performed the experiments, analyzed the data, prepared figures, wrote the paper, reviewed drafts of the paper; KVDW contributed reagents / materials / analysis tools, reviewed drafts of the paper; BJMB conceived and designed the experiments, performed the experiments, contributed reagents / materials / analysis tools, reviewed drafts of the paper.

## Human Ethics

The subjects were informed as to the full nature and design of the study and gave written consent to be participants. This study and its associated research protocols were approved by the Institutional Review Board at the University of Oregon on 2013 December 23 (reference # 03112016.016). All researchers assigned to this protocol were CITI certified to work with human subjects. Identities of participants were never recorded on samples or in resulting datasets.

## Data Deposition

All sequence data and scripts used for statistical analysis have been deposited in the open access data repository Figshare: 10.6084/m9.figshare.7397477

## Supplemental Information

**Supplemental Figure 1:**
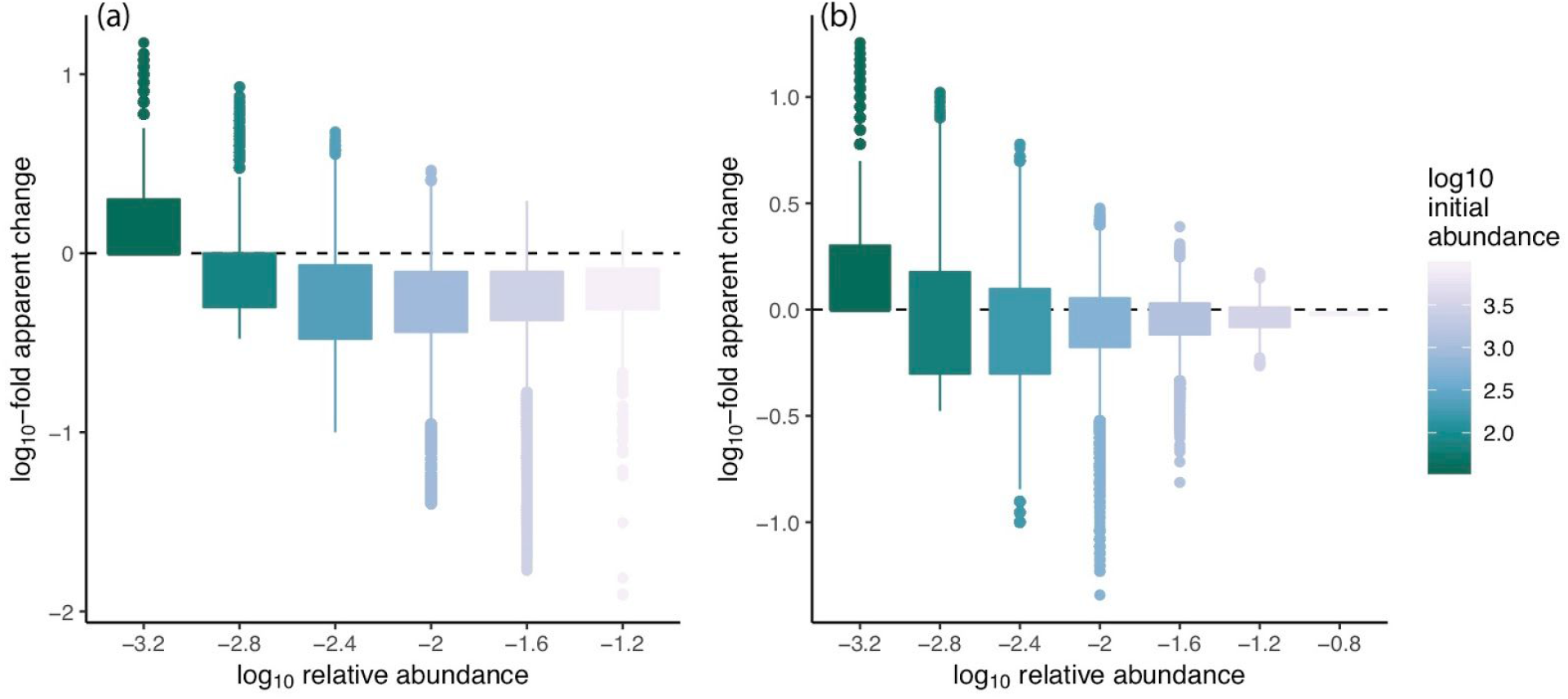
Sampling simulation models (10,000 permutations) as in Fig. 6, but showing (a) the effect of fixed depth sampling (in this case, 1000 sequences per sample) in place of sampling at a fixed percentage of available sequences, and (b) the effect of sampling from a “control transfer” mimicking the control transfers in the empirical experiment, with the source community being the same as the recipient community. Note that the apparent increase in rare taxa is indicated in both alternative simulations.

**Supplemental Figure 2:**
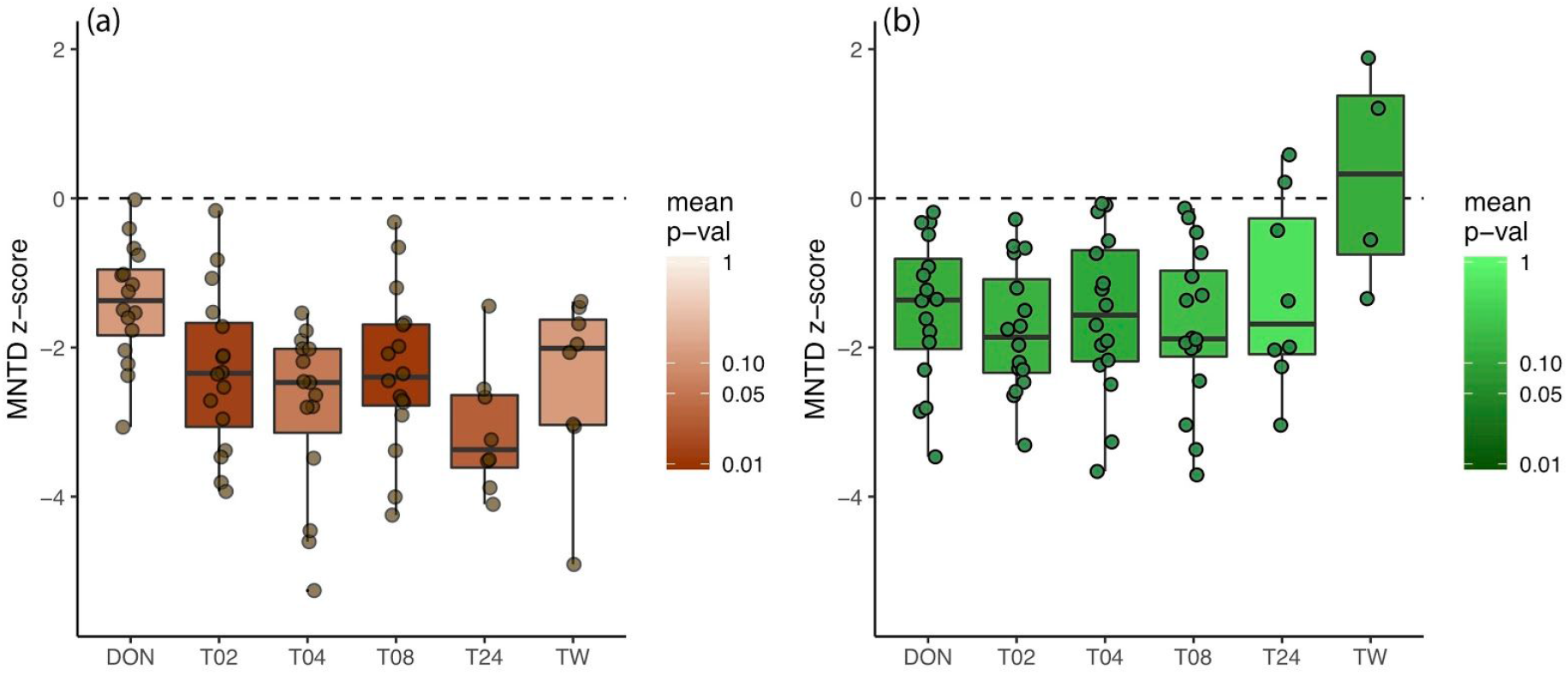
Phylogenetic clustering, measured by the permutational standardized effect size (*z*-score) on mean nearest taxon distances (MNTD) from soil (a) and leaf (b) simulated microbial transfer events.

**Supplemental Figure 3:**
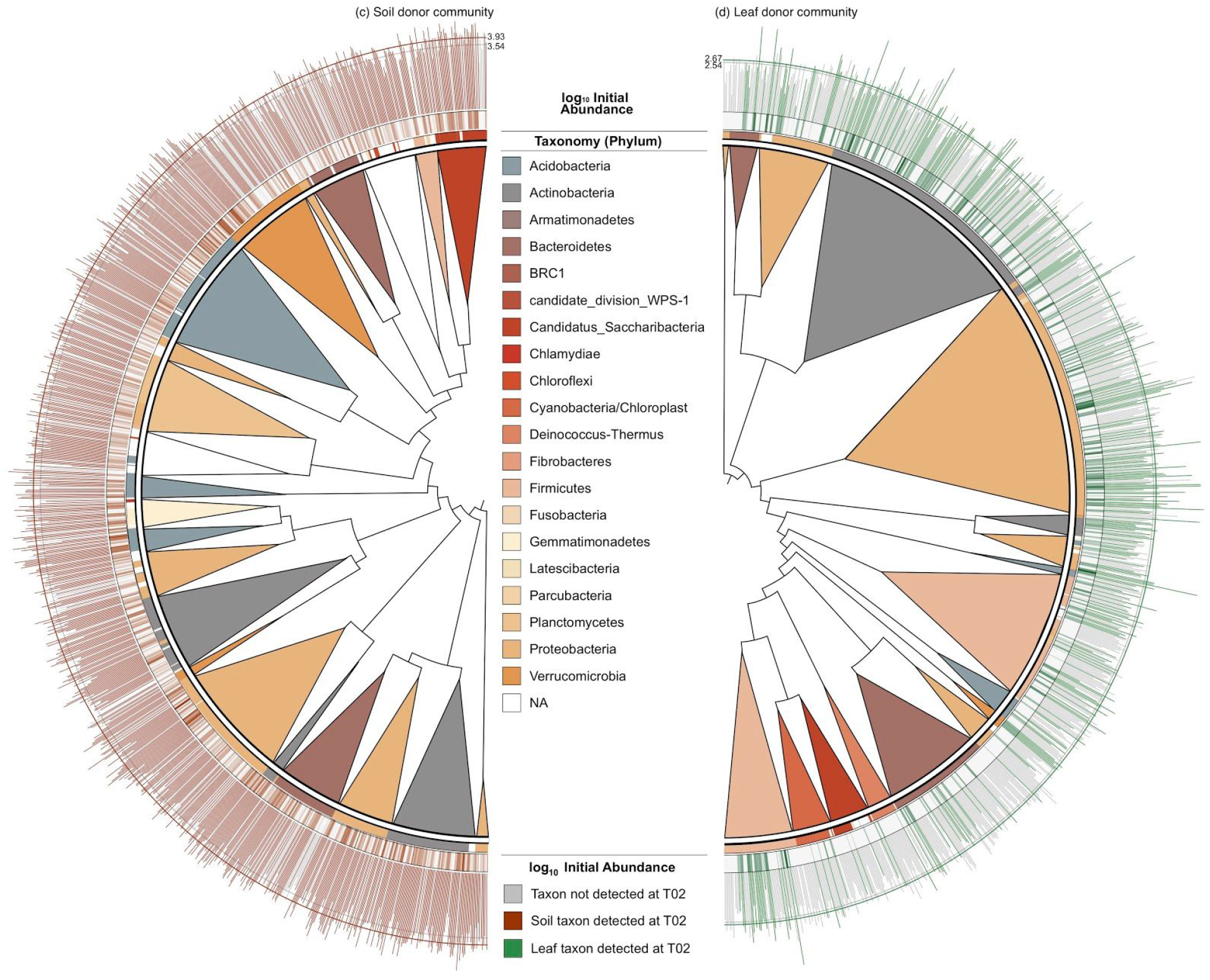
FastTree 16S tree reconstructions for soil (a) and leaf (b) donor microbial communities. Taxa were grouped into polytomies at approximately the phylum level, with the width of the polytomy corresponding to the number of taxa included, the root of the polytomy representing the first branching within the lineage, and the color of the polytomy triangle corresponding to the dominant phylum-level taxonomic assignment for taxa within that lineage. Individual taxonomic assignments are shown in the ring directly above the tree. A heatmap of log_10_-transformed initial absolute abundance is displayed in the ring above the taxonomic assignments. Finally, a bar graph of log_10_-transformed initial absolute abundance is displayed above that, with colored bars indicating those taxa that were detected on the skin after simulated microbial transfer event (at the first sampling, T02); mean transformed abundance for both total initial (grey) and only those taxa that transferred (colored) are indicated by scale rings. The heatmap and the bar graph are presented together to clearly illustrate the effect of abundance on detection post-transfer event.

**Supplemental Figure 4:**
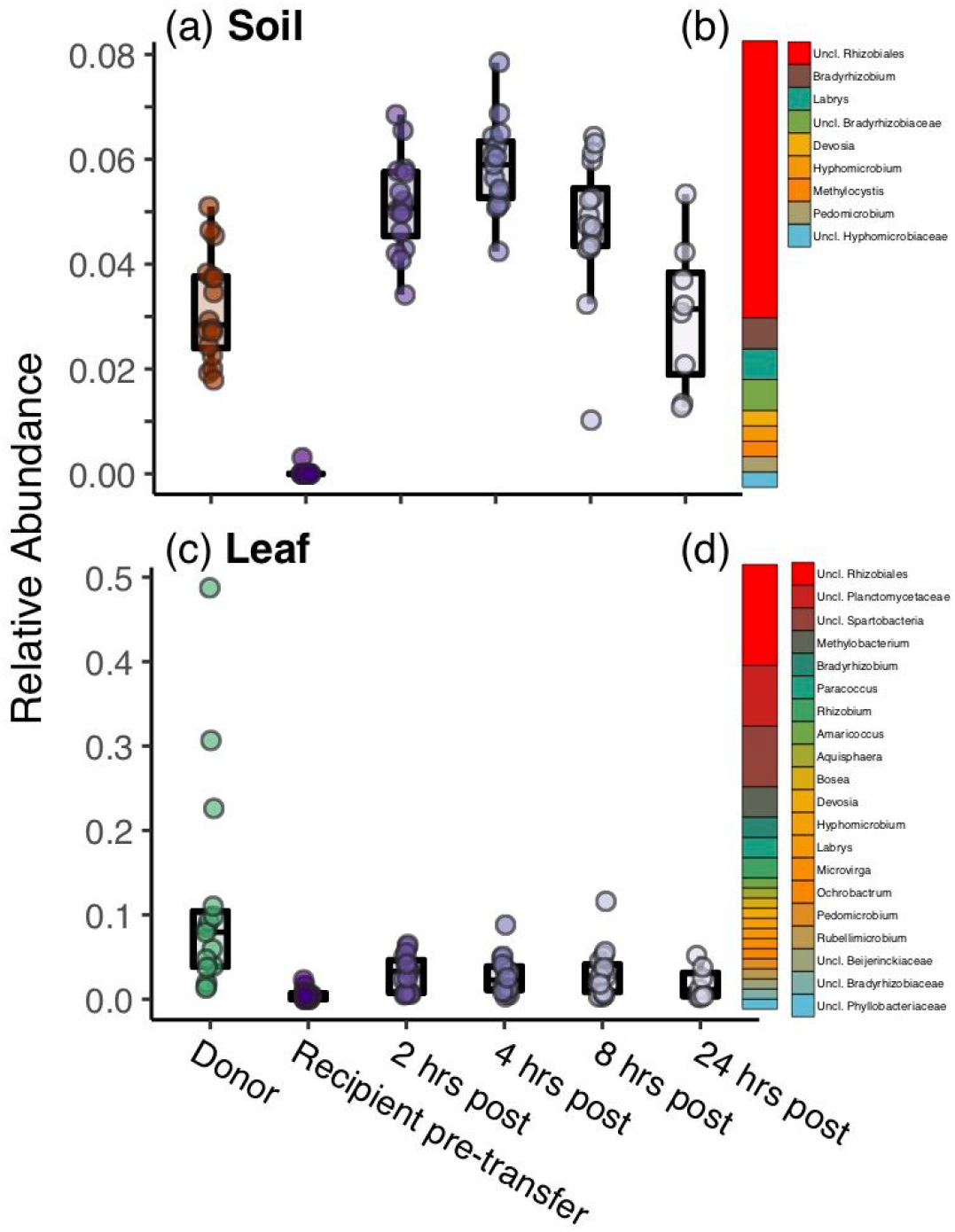
Results from phylogenetic tree-informed sparse discriminant analysis (sLDA). Dynamics for both soil (a) and leaf (c) simulated microbial transfer events are shown for the most discriminant clade selected in each group; the relative composition of each clade is shown (b, soil transfer; d, leaf transfer). A phylogenetically related group of microorganisms including the genus *Bradyrhizobium* (a, b) was selected by the sLDA as indicative of soil microbial transfer; *Bradyrhizobium* are common soil bacteria (94), many of which are nitrogen-fixers, often forming nodules on the roots of legumes (95). The presence and relative abundance of this group, including *Bradyrhizobium*, is strongly indicative of the soil simulated transfer event across subjects. In the leaf simulated transfers, the groups selected by sLDA included a clade in the Alphaproteobacteria (c, d), including members of the Rhizobiales such as *Rhizobia* and *Methylobacterium*, common phyllosphere bacteria which include nitrogen fixers (96) often associated with the leaves of plants (97, 98). These results reinforce the finding from phylogenetic analysis and mixed effects modeling on initial biomass that simulated microbial transfer events lead to the mixing of species abundance distributions, selecting common members of the donor communities and adding them to the recipient community. Numerical dominance of common donor microbes decays with time, allowing typical human microbiome members found in the baseline samples to again be detected in later samples.

